# The conserved protein WhiA influences branched-chain fatty acid precursors in *Bacillus subtilis*

**DOI:** 10.1101/2022.11.02.514974

**Authors:** Laura C. Bohorquez, Joana de Sousa, Transito Garcia-Garcia, Gaurav Dugar, Biwen Wang, Martijs J. Jonker, Marie-Françoise Noirot-Gros, Michael Lalk, Leendert W. Hamoen

**Author notes:** For correspondence: L. W. Hamoen.

## Abstract

The conserved WhiA protein family is present in most Gram-positive bacteria and plays a role in cell division. WhiA contains a DNA-binding motive and has been identified as a transcription factor in actinomycetes. In *Bacillus subtilis*, the absence of WhiA influences cell division and chromosome segregation, however, it is still unclear how WhiA influences these processes, but the protein does not seem to function as transcription factor in this organism. To further investigate the function of WhiA in *B. subtilis*, we performed a yeast two-hybrid screen to find interaction partners, and a Hi-C experiment to reveal possible changes in chromosome conformation. The latter experiment indicated a reduction in short range chromosome interactions, but how this would affect either cell division or chromosome segregation is unclear. Based on adjacent genes, a role in carbon metabolism was put forward. To study this, we measured exometabolome fluxes during growth on different carbon sources. This revealed that in Δ*whiA* cells the pool of branched-chain fatty acid precursors is lower. However, the effect on the membrane fatty acid composition was minimal. Transcriptome data could not link the metabolome effects to gene regulatory differences.

**IMPORTANCE:** WhiA is a conserved DNA binding protein that influences cell division and chromosome segregation in the Gram-positive model bacterium *B. subtilis*. The molecular function of WhiA is still unclear, but a previous study has suggested that the protein does not function as a transcription factor. In this study, we used yeast two-hybrid screening, chromosome conformation capture analysis, metabolomics, transcriptomics and fatty acid analysis to obtain more information about the workings of this enigmatic protein.

## INTRODUCTION

WhiA is a conserved DNA binding protein that can be found in most Gram-positive bacteria, including the simple cell wall-lacking *Mycoplasmas*. The crystal structure of *Thermotoga maritima* WhiA shows a bipartite conformation in which a degenerate N-terminal LAGLIDADG homing endonuclease domain is tethered to a C-terminal helix-turn-helix DNA binding domain. However, none of the characterized WhiA proteins have shown any nuclease activity (1). In the actinomycetes *Streptomyces*, *S. venezuelae* and *Corynebacterium glutamicum* WhiA functions as a transcriptional activator of many genes, among which the key cell division gene *ftsZ* (2–4). Mutations in *whiA* prevent the induction of FtsZ in streptomycetes, thereby blocking synthesis of sporulation septa (5, 6). In *Bacillus subtilis*, inactivation of WhiA reduces the growth rate in rich medium and affects the expression of a variety of genes, but not that of *ftsZ* or other cell division related genes (7). Moreover, no relationship was found between WhiA binding sites on the genome and regulated genes, suggesting that WhiA does not function as a classic transcription factor in this organism (7). Nevertheless, WhiA is important for cell division in *B. subtilis*, and the absence of WhiA is synthetic lethal when cell division proteins are inactivated that regulate the formation of the Z-ring, such as the regulatory MinCD proteins, and the FtsZ polymer crosslinker ZapA (7). Later it was found that WhiA is also important for proper chromosome segregation in this organism, and *whiA* mutants display increased nucleoid spacing (8). Despite the conserved nature of this protein and its role in key cellular processes, it is unclear how this protein operates in *B. subtilis*. In the current study, we performed a wide variety of analysis, including a Yeast two-hybrid analysis, chromosome conformation capture (Hi-C), metabolomics and transcriptomics, to gain a better understanding of the function of WhiA. Eventually, this led us to investigate the fatty acid composition of the cell membrane in *whiA* mutants.

## RESULTS

### Yeast two-hybrid screening

To find possible interaction partners of WhiA that could help to elucidate its function, we performed a genome wide yeast two-hybrid screen (9). To increase the chances of detecting relevant interactions, we used full length WhiA, and separately its N- and C-terminal domains, containing the degenerative LAGLIDADG homing endonuclease domain (amino acids 1-227) and the helix-turn-helix domain (amino acids 222-316), respectively. The latter domain is responsible for interaction with the chromosome, which was confirmed by a microscopic analysis of GFP fusions (Fig. S1). After screening a genomic library with an approximately 15-fold redundancy of the *B. subtilis* genome, we found 3 potential interaction partners, YlxS, YrhJ and YlaD, which interacted both with full length WhiA and the N-terminal domain (Fig. S2A). Full-length WhiA showed some auto-activation in the screen. When we used the synthetic complete media lacking leucine, uracyl and adenine (-LUA), which makes the selection more stringent (9), the interaction between YlxS and full length WhiA was still observed (Fig. S2A). YlxS is 32 % identical to *Escherichia coli* RimP involved in ribosome assembly (10). YrhJ is a fatty acid monooxygenase, catalyzing hydroxylation of a range of fatty acids (11), and YlaD functions as an anti-sigma factor (12, 13). To test whether these proteins are involved in the activity of WhiA, we deleted the corresponding genes and tested for reduced growth rate in rich medium, chromosome segregation defects, and cell division phenotype in a Δ*zapA* background strain. Unfortunately, none of the deletion mutants showed a phenotype that resembled that of a Δ*whiA* mutant (Fig. S2B). It is therefore unlikely that either YlxS, YrhJ or YlaD is involved in the activity of WhiA.

### Chromosome conformation

A conserved feature of Δ*whiA* mutants is the increased internucleoid distance (8). Since WhiA is a conserved DNA binding protein it might play a role in the organization of the chromosome. To examine this, we performed a Hi-C (chromosome conformation capture) analysis of Δ*whiA* cells. Both wild type and Δ*whiA* cells produced similar contact maps and the absence of WhiA does not affect the alignment of chromosome arms by the SMC condensing complex (Fig. 1A) (14). However, a clear difference was observed for short range genome interaction between the two strains (Fig. 1B). Short range interactions (< 50 kb) were reduced upon *whiA* deletion, thereby indicating potential involvement of WhiA in mediating non-specific local interaction on a genome wide scale. However, it is unclear how this would lead to increased spacing between daughter chromosomes or influence cell division.

**Fig. 1.**
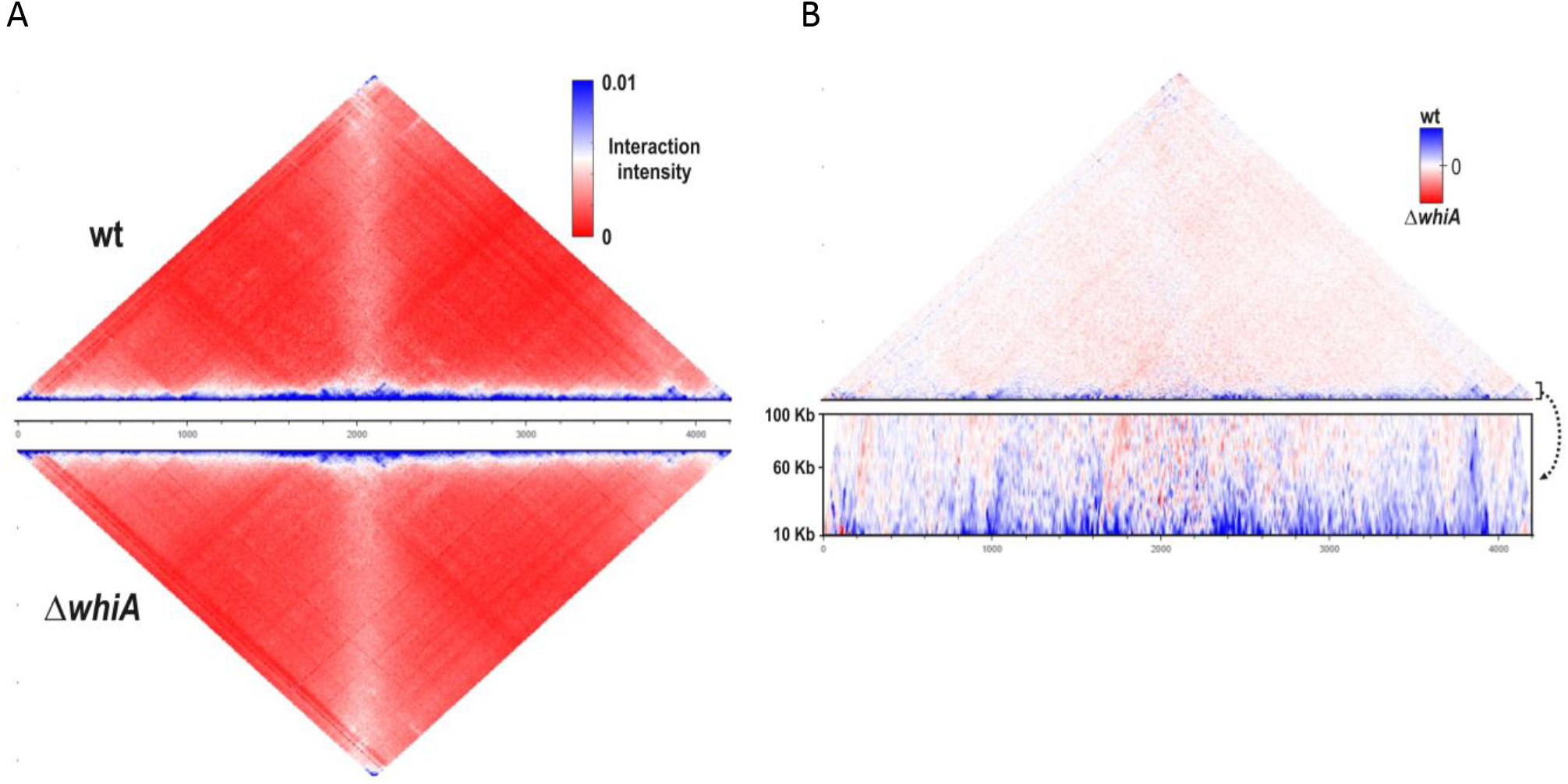
Chromosome conformation capture (Hi-C) analysis. (A) Normalized Hi-C contact maps of wild type (top) and Δ*whiA* strains (below) at exponential phase. SMC dependent juxtaposition of the chromosome arms is observed in both strains as the secondary (vertical) diagonal (14). (B) Difference plot of wild type and Δ*whiA* strains. The magnified view of difference in short-range contacts (between 10 kb and 100 kb) is shown below.

### Growth on different carbon sources

*whiA* is the 4^th^ gene in an operon of 6 genes that is constitutively expressed during growth (Fig. S3) (7). The first gene, *yvcI*, encodes a Nudix hydrolase that hydrolyses organic pyrophosphates and is considered a housecleaning enzyme (15, 16). The second gene, *yvcJ*, encodes a GTPase required for the proper expression of DNA uptake proteins during natural competence (17, 18). The third gene, *yvcK*, encodes an UDP-sugar binding protein that is essential for growth under gluconeogenic conditions (19). *crH*, the fifth gene downstream of *whiA*, is a HPr-like protein that participates in catabolite repression as secondary cofactor of the global catabolite regulator CcpA (20, 21). The final gene, *yvcN*, is an uncharacterized acetyltransferase (Subtiwiki database (16)). In many bacteria *whiA* is located adjacent to *yvcK* and *crh* (STRING database (22)). Possibly, this conserved organization points towards a metabolic function of WhiA. Inactivation of *yvcK* blocks growth on citrate and result in very poor growth on either fumarate or malate as sole carbon sources (23). To examine whether the absence of WhiA also affects growth using these carbon sources, we grew a *whiA* mutant in Spizizen minimal salt medium using either malate, fumarate or citrate as carbon source. To prevent any downstream effects, a marker-less *whiA* mutant was used, containing a stop codon at the beginning of the gene (strain KS696 (7)). As shown in Fig. 2A, the *whiA* mutant was able to grow in the different media with a growth rate similar to that of the wild-type strain, indicating that WhiA and YvcK work in different pathways. As shown in Fig. 2A and previously reported, the *whiA* mutant grows slower in rich LB medium (7). The reason that this effect was not observed in minimal medium (Fig. 2A), could be related to the lower doubling time in minimal medium compared to LB (~53 min versus ~21 min), which can mitigate chromosome segregation and cell division defects (24, 25). Therefore, we tested whether the chromosome segregation and cell division defects were present in minimal medium. Interestingly, the inter-nucleoid spacing was still larger in a *whiA* mutant (Fig. S4A), and depletion of WhiA in a Δ*zapA* background increased the cell length and occasionally generated aberrant nucleoids in minimal medium (Fig. S4B).

**Fig. 2.**
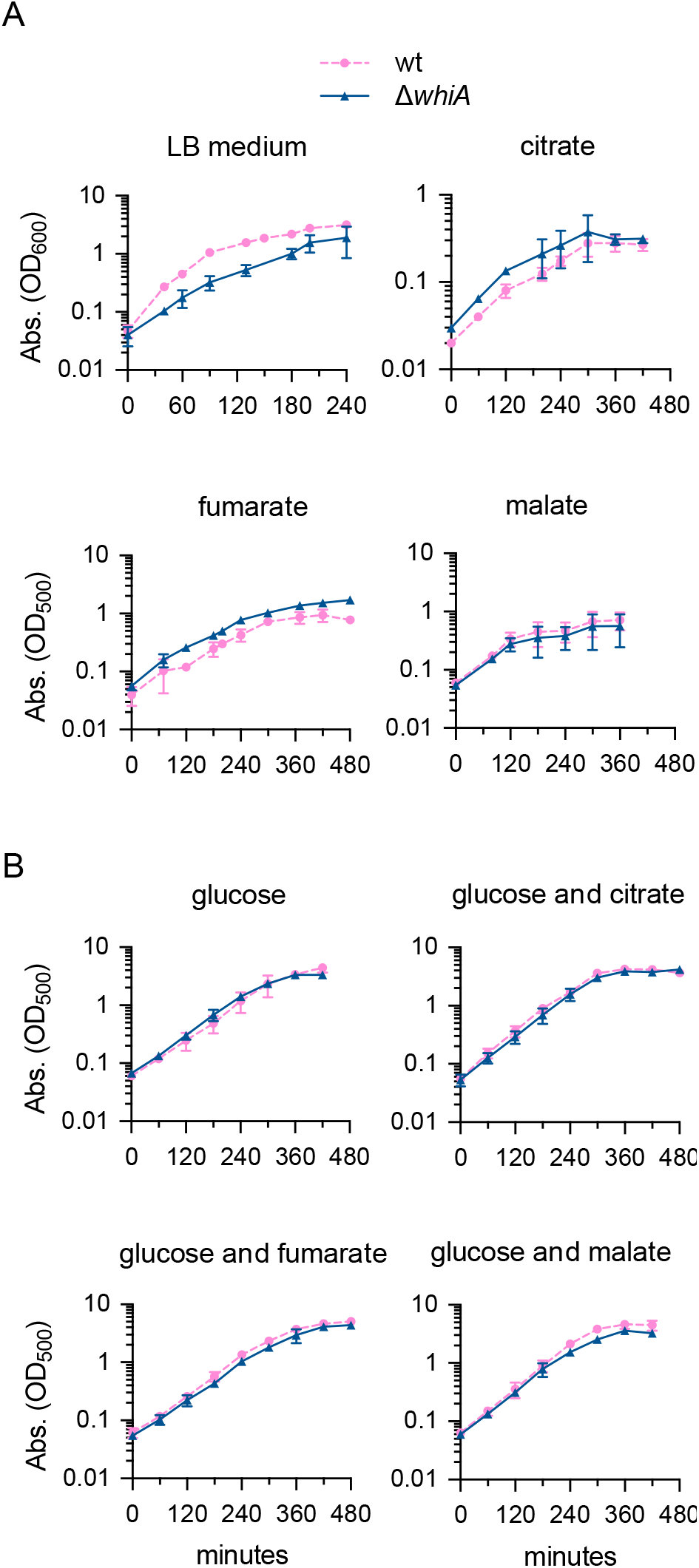
Growth on different carbon sources. Growth measured as optical density of the wild-type strain (strain 168) and the *whiA* marker-less mutant (strain KS696). (A) Growth in LB medium, and in Spizizen minimal salt medium (SMM) supplemented with 22 mM of either citrate, fumarate or malate. (B) Growth in chemically defined minimal Amber medium supplemented with 22 mM of either glucose, glucose and citrate, glucose and fumarate or glucose and malate. Data are shown as mean values and standard deviation of triplicate samples.

### Utilization of carbon sources

To examine whether WhiA is involved in catabolite regulation, like *crh*, we first measured the carbon consumption by means of exometabolomics, using proton nuclear magnetic resonance spectroscopy (^1^H-NMR) (26). This required a minimal chemically defined medium for which often M9 medium is used. However, M9 medium has been optimized for *E. coli* and not for *B. subtilis*, and the latter easily lyses in this medium in the stationary phase (26). Therefore, we composed an alternative chemically defined medium based on different minimal media used for *B. subtilis*, as listed in Table S1. In essence, the resulting medium, named Amber medium, uses a phosphate buffer, ammonium salt and glutamate as nitrogen sources, and 22 mM for any carbon source. We tested growth on glucose alone, glucose and citrate, glucose and fumarate, and glucose and malate. Fig. 2B shows that both wild type and the marker-less *whiA* mutant grows fine in this medium using these conditions. Malate was incorporated in this analysis since it is the second preferred carbon source of *B. subtilis*, and its utilization is not subjected to carbon catabolite repression in this organism (27).

To determine the exometabolome, 2 ml of culture was collected at regular time intervals and rapidly filtered, and the filtrate stored at −20 °C for later ^1^H-NMR spectroscopic analysis. Identification of metabolites was based on NMR spectra alignment of pure standard compounds and the quantification was done based on the integration and comparison of the designated peaks to an internal standard peak (see methods section for details). The final data were based on 3 independent biological replicates, and the quality of the replicates was confirmed using a principal component analysis (Fig. S5). As shown in Fig. 3, the consumption of the different carbon sources was the same for wild-type and *whiA* mutant cells in all 4 growth conditions. Citrate and fumarate utilization was initiated when most glucose was exhausted, confirming that fumarate and citrate were subjected to glucose-dependent catabolite repression in both strains. Malate was consumed faster than glucose as has been shown before (Meyer *et al*., 2014). These data show that WhiA is not involved in catabolite repression.

**Fig. 3.**
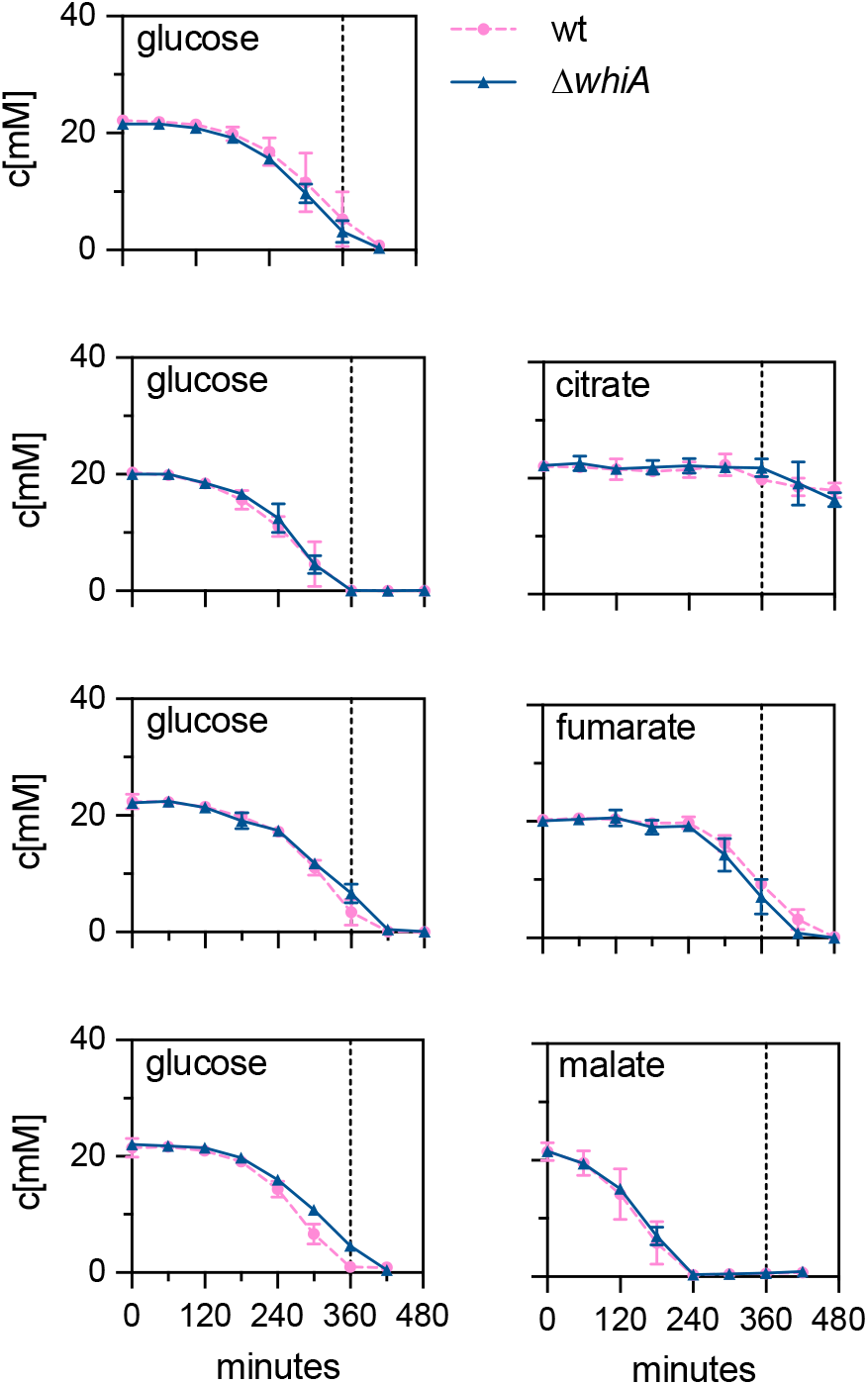
Carbon utilization. Carbon source utilization (concentration in mM) of wild-type (strain 168) and *whiA* marker-less mutant cells (strain KS696) during growth in defined minimal (Amber) medium supplemented with either glucose, glucose and malate, glucose and citrate or glucose and fumarate (22 mM each). Data are shown as mean values and standard deviation of triplicate samples. The dashed lines mark the time point when the glucose culture enters stationary phase (360 min) (see Fig. 2B).

### Exometabolome analysis

Aside of the supplied carbon sources (glucose, citrate, fumarate and malate), we were able to detect 18 other metabolites in the medium. Interestingly, several of these metabolites showed a different secretion kinetic in the *whiA* mutant. To facilitate the interpretation of the exometabolome data, the time-resolved extracellular metabolite concentrations were plotted onto the relevant pathways (Fig. 4 and 5). The differences became apparent after approximately 180 min, when glucose levels started to go down. The depleted pools of the branched-chain fatty acid precursors isovalerate, isobutyrate and 2-methylbutyrate (Fig. S7A), and the higher secretion of acetate and 2-oxoglutarate in the *whiA* mutant, are most obvious. We were not able to identify isoleucine, leucine and oxaloacetate due to the detection limits of the method (26). Citrate and isocitrate were only measurable when the medium contained the TCA intermediate citrate or fumarate (Fig. 4 lower panel, and Fig. 5 upper panel). The reason for this is that expression of citrate synthase and aconitase is induced when citrate is present in the medium or fumarate becomes the sole carbon source after glucose levels have fallen (26, 28).

**Fig. 4.**
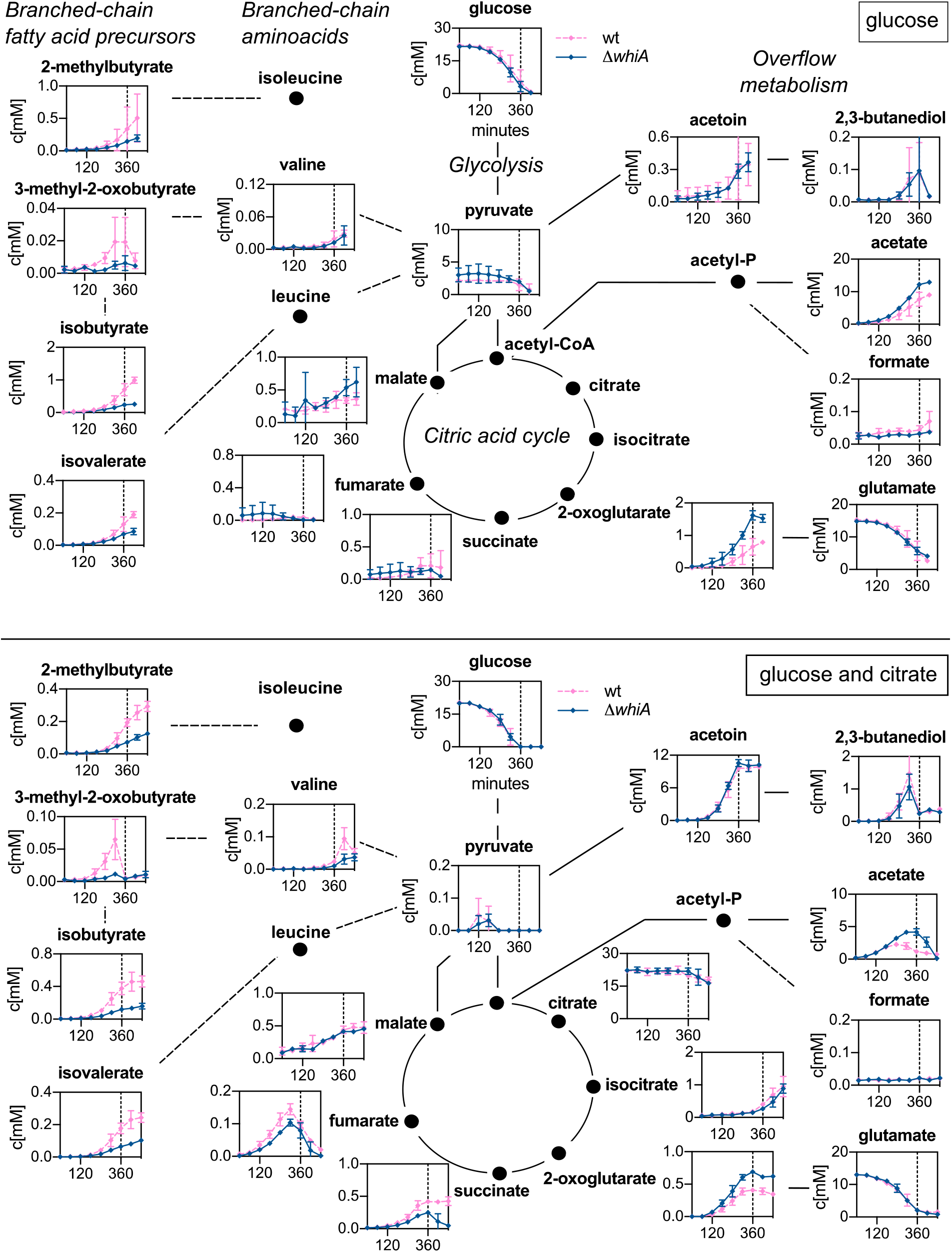
Exometabolome of cells grown with either glucose or glucose and citrate. Time-resolved extracellular metabolite concentrations (in mM) of wild-type (strain 168) and *whiA* marker-less mutant cells (strain KS696) grown in chemically defined minimal Amber medium with either glucose alone (upper panel) or glucose and citrate (lower panel) as carbon source (22 mM each). Dashed lines indicate entry into stationary phase (360 min). The compounds are arranged according to the main metabolic pathways: glycolysis, TCA cycle, overflow metabolites, branched-chain amino acids and branched-chain fatty acids precursors. Data are shown as mean values and standard deviation of triplicate samples.

**Fig. 5.**
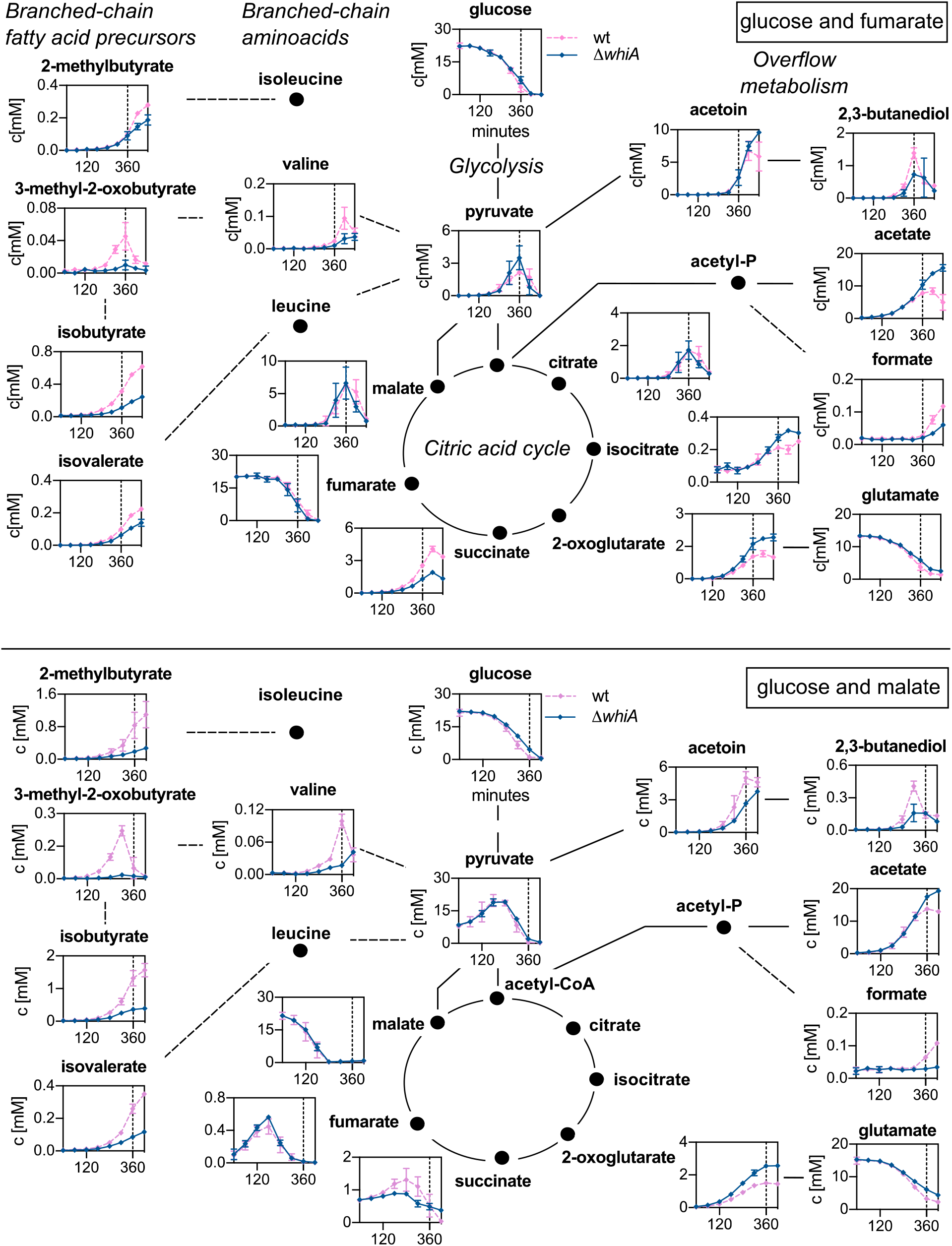
Exometabolome in cells grown with either glucose and fumarate or glucose and malate. Time-resolved extracellular metabolite concentrations (in mM) of wild-type (strain 168) and *whiA* marker-less mutant cells (strain KS696) grown in chemically defined minimal Amber medium with either glucose and fumarate (upper panel) or glucose and malate (lower panel) as carbon sources (22 mM each). Dashed lines indicate entry into stationary phase (360 min). The compounds are arranged according to the main metabolic pathways: glycolysis, TCA cycle, overflow metabolites, branched-chain amino acids and branched-chain fatty acids precursors. Data are shown as mean values and standard deviation of triplicate samples.

### Transcriptome analysis

To examine whether the changes in metabolism were related to changes in gene expression, we compared the transcriptomes of wild type and *whiA* mutant cells grown in Amber medium supplemented with glucose and malate as carbon sources. When the cultures reached an OD_500_ of 0.5 (Fig. 2B, 120-180 min), cells were harvested for RNA isolation. The experiment was repeated one more time to provide a biological replicate. The volcano plot in Fig. 6 depicts the distribution of expression differences against adjusted *p*-values. 57 genes were upregulated and 40 downregulated more than 3-fold with an adjusted *p*-value < 0.05 (Table 1, data for all genes are listed in Table S7). The most highly upregulated genes, *ydcF, ydcG* and *pamR* form an operon. PamR is a transcription factor that affects expression of prophages and certain metabolic genes (29). The *bmrB* operon, coding for a multidrug ABC transporter (30), is also strongly upregulated in the *whiA* mutant. This transporter is involved in the activation of KinA, one of the key regulators of sporulation. It should be mentioned that a *whiA* mutant displays only a very mild defect in sporulation (7). The upregulated *tapA* operon is required for synthesis of the major extracellular matrix (31). Other upregulated genes were the *epeX (yydF*) operon encoding proteins controlling the activity of the LiaRS cell envelope stress-response system (32), the *fatR* operon involved in lipid degradation (33), and the*, sunA* and *nupN operons* necessary for biosynthesis of a siderophore, antimicrobial peptide and the uptake of guanosine, respectively (*34–36*).

**Fig. 6.**
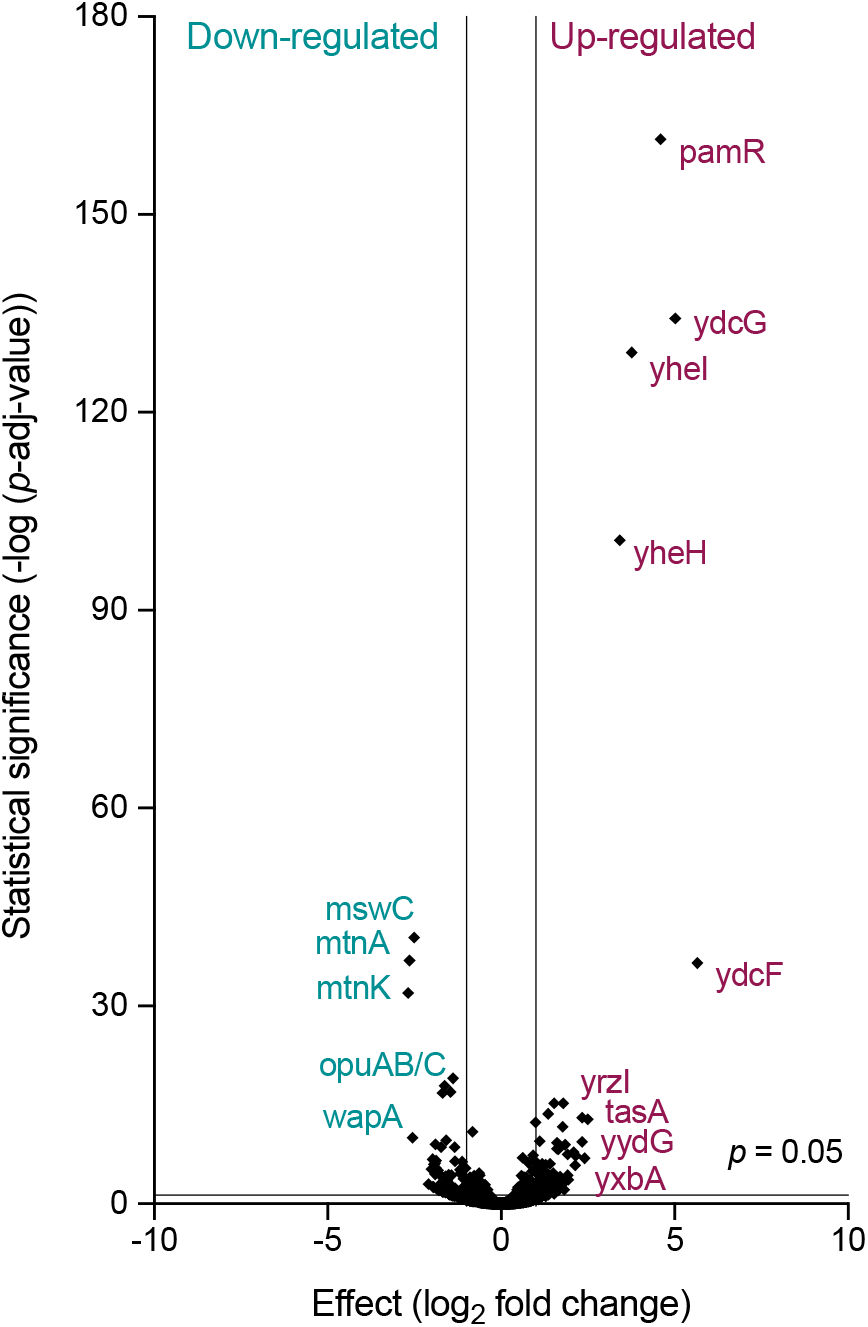
Volcano plot of transcriptome data. Volcano plot depicting the transcriptome data as a relation between adjusted *p*-values and log_2_ fold expression change. Wild-type (strain 168) and *whiA* marker-less mutant cells (strain KS696) were grown in defined minimal (Amber) medium with glucose and malate and sampled during exponential growth. Main downregulated and upregulated genes in the *whiA* mutant are shown in green and red, respectively. Genes are listed in Table S2.

**Table 1.**
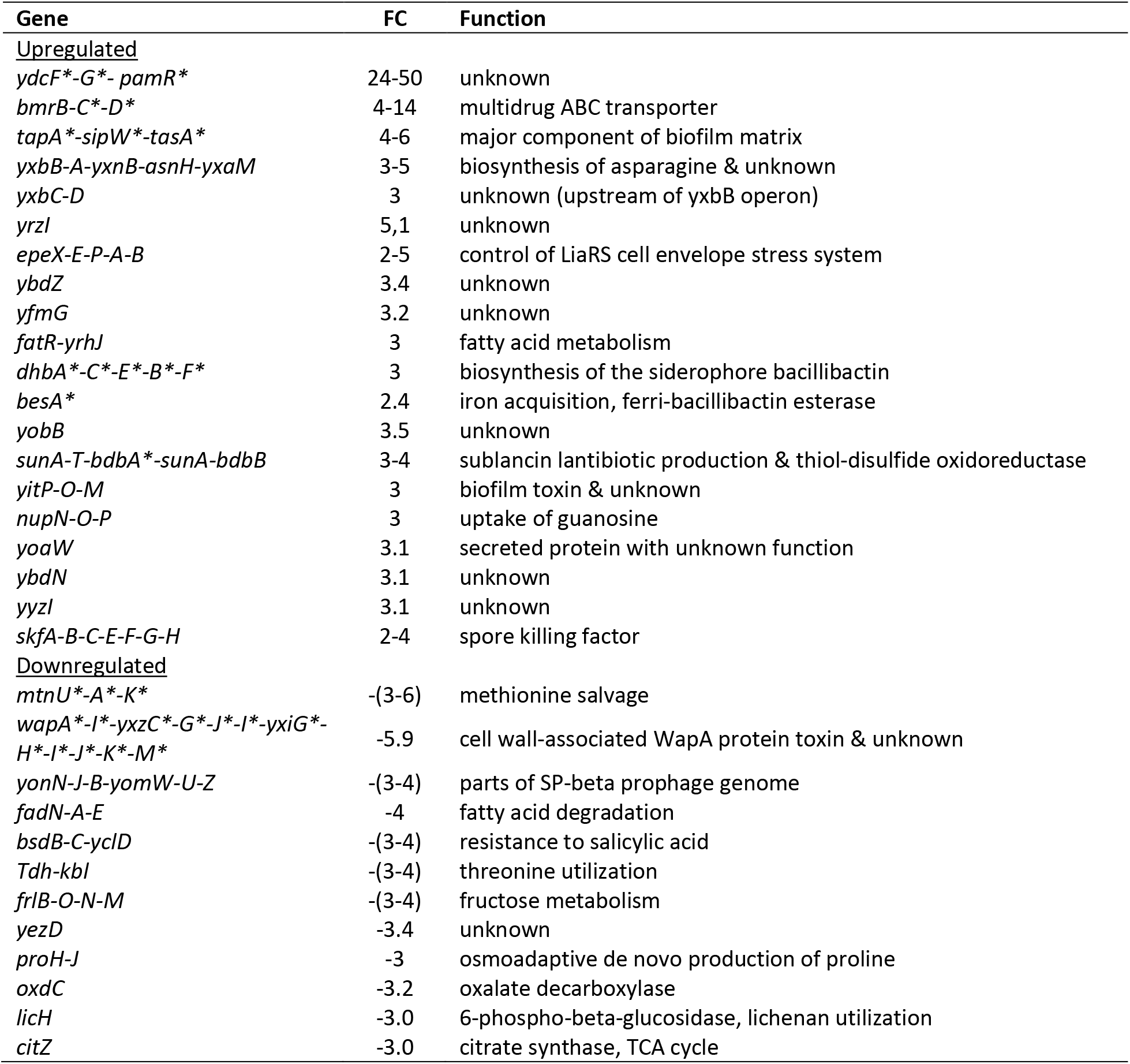
Transcriptome comparison of wild-type (strain 168) and *whiA* marker-less mutant cells (strain KS696). Cells were grown in defined minimal (Amber) medium with glucose and malate and harvested for RNA isolation during exponential growth (OD_600_ ~0.5). Genes with an adjusted *p*-value < 0.05 and Fold Change (FC) > 3 (Δ*whiA*/wt) are listed. Genes found in a previous transcriptome Δ*whiA* analysis performed in LB rich medium are indicated by * (7) (see also Fig. S8). Genes located in one operon are listed together in one row.

Strongly downregulated genes comprised the *wapA* operon, expressing one of the main cell surface proteins in *B. subtilis* (37), the *fadN* operon involved in fatty acid degradation (38), and the *frlB* operon coding for an amino sugar uptake system (39). Several genes involved in amino acid biosynthesis were also downregulated, including the *mtnA* operon involved in methionine salvage (40), the *tdh* operon involved in threonine utilization (41, 42), and *proHJ* necessary for production of proline (43), respectively. Finally, expression of the major citrate synthase encoded by *citZ* was also significantly downregulated (44).

The downregulation of citrate synthase did not show in the exometabolomics data, and in fact the secretion of 2-oxoglutarate, downstream of citrate in the TCA cycle, was higher in the *whiA* mutant (Fig. 5). In the medium containing citrate or fumarate as additional carbon sources, there was also no difference in either citrate or isocitrate secretion and consumption between wild type and the mutant (Fig. 4 and 5). When we lowered the stringency and included genes that were more than 2-fold regulated (Table S7), only the 2.8-fold upregulation of *rocG*, which encodes glutamate dehydrogenase responsible for the conversion of glutamate to 2-oxoglutarate (45), could be linked to the metabolomics data, since 2-oxoglutarate levels increased faster in the *whiA* mutant in all four growth conditions (Fig. 4 and 5). However, another reason for the increased 2-oxoglutarate levels might be the reduction in branched-chain fatty acid precursors that rely on 2-oxoglutarate for the aminotransferase reaction (46). The transcriptome data did not reveal any obvious reason for the reduced synthesis of branched chain fatty acid precursors. Table S2 lists the fold-change expression of the main genes involved in branched-chain amino acids metabolism and fatty acid synthesis (see Fig. S7 for pathway schemes). The branched-chain amino acid transporters *bcaP* and *braB* were upregulated significantly by 1.9 and 1.4-fold, respectively (*p*-value<0.05), and so were *ybgE* and *ilvD* involved in branched-chain fatty acid precursors synthesis (1.9- and 1.5-fold, respectively). Possibly, this is a response to low substrate levels. However, *yvbW*, encoding a putative leucine permease, was downregulated 1,7-fold. The *leuA* operon involved in leucine biosynthesis was downregulated significantly but only by approximately 1.4-fold, and there was no significant difference in expression of either valine or isoleucine biosynthesis genes (Table S2). Overall, the transcriptome data did not provide a clear explanation for the exometabolome differences.

### Fatty acid analysis

*B. subtilis* contains primarily branched-chain fatty acids. Synthesis of anteiso-fatty acids requires isoleucine, and the iso-C15 and -C17 and iso-C14 and -C16 fatty acids require leucine and valine, respectively (de Mendoza *et al*., 2002). Therefore, the reduced cellular concentration of these amino acids might lead to a change in the fatty acid composition of a *whiA* mutant. To investigate this, we analyzed the fatty acid composition of wild-type (strain 168) and Δ*whiA* cells (strain KS696) using gas chromatography (Table S3). For this, cells were harvested when the cultures reached an OD_500_ of approximately 0.5. The majority of fatty acids, 93.9 % in the wild type strain, are branched-chain fatty acids, and this fraction hardly changes in the Δ*whiA* mutant (93.1 %). The distribution of straight, iso and anteiso chains over the different fatty acids is shown in Fig. 7A. The fraction of iso-fatty acids in the *whiA* mutant is slightly down from 53.8 % to 45.8 %, whereas the fraction of anteiso fatty acids slightly increases from 40.1 % to 47.3 % (Fig. 7B). The reduction in leucine and valine derived fatty acids is in line with the metabolome data, but the increased contribution of isoleucine derived fatty acids is not.

**Fig. 7.**
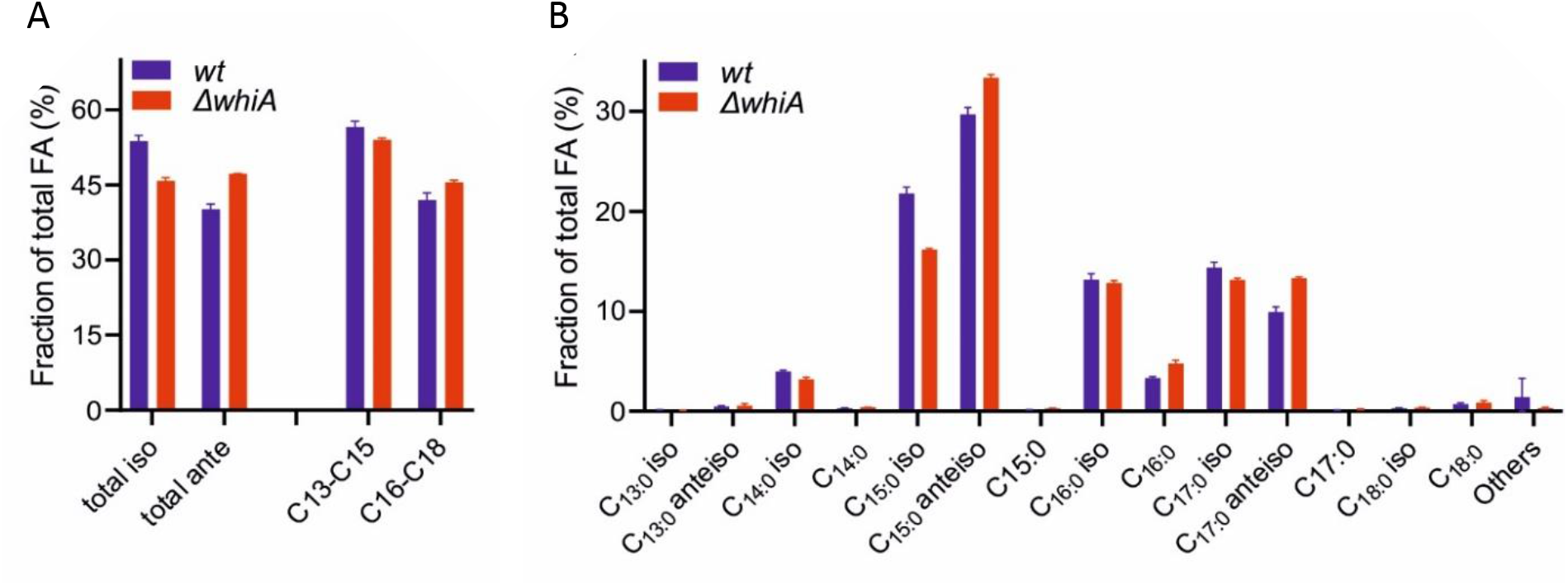
Fatty acid analysis of Δ*whiA* mutant. (A) Comparison of the total iso- and anteiso-fatty acids and fatty acid chain length between wild type *B. subtilis* and Δ*whiA* cells. (B) Detailed comparison of the different fatty acids. Concentrations of individual fatty acids are listed in Table S3.

Anteiso-fatty acids disturb the lipid packing more than iso-fatty acids and will therefore increase membrane fluidity, which is an important way *B. subtilis* regulates its membrane fluidity (47). This might explain why the Δ*whiA* mutant contains 2.5 % less short fatty acid species (C13, C14, C15) and 3.6 % more long fatty acid species (C16, C17, C18) (Fig. 7A), in order to maintain membrane fluidity homeostasis. Indeed, a membrane fluidity assay, using the membrane fluidity sensitive dye Laurdan (48, 49), did not detect strong differences in membrane fluidities between both strains when grown in either minimal medium or LB (not shown).

## DISCUSSION

Despite the conserved nature of WhiA and its documented role as transcriptional activator in the actinomycetes, it is unclear how this protein functions in *B. subtilis*, and other Gram-positive bacteria. Hi-C data indicated that the absence of WhiA reduces short range (< 50 kb) chromosome interactions. The areas where this occurred did not correlate to WhiA binding sites that were previously determined using Chip-on-chip analysis (7). We could also not detect a clear correlation between transcription difference and the absence of these short range chromosome interactions (data not shown). However, the current resolution of the Hi-C analysis is insufficient to make such correlations. Whether the reduction in short range chromosome interactions affects chromosome segregation is unclear since there is no clear mechanism that would link these two phenomena.

Both cell division and chromosome replication are linked to the metabolic state of cells. In *B. subtilis* the glycosyltransferase UgtP couples nutritional availability to cell division (50). The protein is involved in lipoteichoic acid synthesis using UDP-glucose as substrate. Under nutrient-rich conditions, intracellular levels of UDP-glucose are high and UgtP inhibits FtsZ assembly in a UDP-glucose dependent manner. Another example is the glycolytic enzyme pyruvate dehydrogenase kinase that, by an yet unknown mechanism, positively regulates Z-ring assembly (51). Moreover, temperature sensitive mutants in the *B. subtilis* DnaC helicase, DnaG primase and DnaE polymerase can be suppressed by mutations in different glycolytic enzymes, among which pyruvate kinase (52). Since inactivation of WhiA affects both cell division and DNA segregation and since *whiA* is located in an operon adjacent to *yvcK* and *crh*, which are involved in gluconeogenic growth and catabolite repression, respectively (19, 20), it was tempting to assume that WhiA affects cell division and chromosome replication by playing a role in carbon metabolism. However, our study showed that WhiA is neither required for gluconeogenic growth nor plays a role in carbon catabolite repression. Nevertheless, the metabolomics data did reveal that in a *whiA* mutant the pool of branched-chain fatty acid precursors is reduced. It was not possible to link this effect to changes in the transcriptome although this might be due to the fact that we measured gene regulation at the end of exponential growth, whereas the differences in the exometabolome was most apparent in the beginning of the stationary phase. Of note, several of the most strongly up- and downregulated genes were also found in a previous transcriptome study where a *whiA* deletion mutant was grown in rich LB medium, including the upregulated *ydcF, bmrB, tasA*, and *dhbE* operons, and the downregulated *mtnK* and *wapA* operons (Fig. S8) (7). However, none of these genes have so far been linked to either cell division or chromosome segregation.

Why the absence of WhiA affects branched-chain fatty acid precursor levels is unclear. It is possible that these changes have an effect on metabolic regulators that use these cofactors, such as CodY, which activity is affected by branched-chain amino acids (53). However, only a small fraction (11 %) of the CodY regulon was significantly affected in the Δ*whiA* mutant. It is also unclear how a reduction in branched-chain fatty acid precursors would influence cell division and DNA segregation in a Δ*whiA* mutant. Changes in the fatty acid composition of the membrane could in theory influence the activity of membrane proteins, however the observed differences were limited, and in fact the addition of the branched-chain fatty acid precursors isovalerate, isobutyrate and 2-methyl butyrate to LB medium did not restore the growth defect of a Δ*whiA* mutant (not shown). In conclusion, the molecular function of WhiA in *B. subtilis*, and therefore in many other Gram-positive bacteria and the mycoplasmas, remains an enigma.

## MATERIALS AND METHODS

### Bacterial strains and growth conditions

Luria-Bertani (LB) medium was used for routine selection and maintenance of *B. subtilis* and *E. coli* strains. Spizizen’s minimal medium SMM (54) consisted of 2 g/l (NH_4_)_2_SO_4_, 14 g/l K_2_HPO_4_, 6 g/l KH_2_PO_4_, 1 g/l sodium citrate, 2 g/l MgSO_4_, 5 g/l glucose, 2 g/l tryptophan, 0.2 g/l casamino acids and 2.2 g/l ammonium ferric citrate. The defined minimal (Amber) medium consisted of 70 mM K_2_HPO_4_ and 30 mM KH_2_PO_4_ (adjusted to pH 7.4), 15 mM sodium chloride, 10 mM (NH_4_)_2_SO_4_, 0.002 mM of trace elements (ZnCl_2_, MnSO_4_, CuCl_2_, CoCl_2_ and Na_2_MoO_4_), 22 mM glucose, 0.25 mM tryptophan, 10 mM glutamate, 1 mM MgSO_4_, 0.1 mM calcium chloride and 0.01 mM ammonium ferric citrate (Table S1). When indicated, the medium was supplemented with 22 mM final concentration of malate, fumarate or citrate. All strains were grown at 37 °C at 250 rpm. *B. subtilis* strains used in this study are listed in Table S4. The mutant strains provided by other labs were transformed into our laboratory strain to ensure isogenic backgrounds. If indicated, the medium was supplemented with a mixture of 3 branched-chain fatty acid precursors (100 μm of 2-methyl-butyrate, isobutyrate and isovalerate, Sigma-Aldrich) or straight fatty acid precursors (100 μm of methyl-butyrate, methyl-propionate and methyl-valerate, Sigma-Aldrich).

WhiA depletion strain (LB45) (8) was always grown in presence of erythromycin, due to the Campbell type integration of the *Pspac-whiA* construct into the *whiA* locus. Depletion of WhiA was accomplished by inoculating a single colony into LB medium with 0.1 mM IPTG and growth at 37 °C to an OD_600_ of ~1. Subsequently, cells were harvested, washed in pre-warmed LB medium, and resuspended to an OD_600_ of 0.01 and grown in the absence of IPTG. For spot dilution assays a single colony (strain LB45) was used to inoculate LB or Amber medium with 0.1 mM IPTG and grown at 37 °C to an OD_600_ of 0.5. Subsequently, cells were serial diluted in pre-warmed LB or Amber medium and 10 μl spots were inoculated and grown at 37 °C overnight.

### Strain constructions

Molecular cloning, PCRs and transformations were carried out using standard techniques. Plasmids and oligonucleotides used in this study are listed in Table S5 and S6, respectively. The xylose inducible msfGFP-WhiA N-terminus, C-terminus and full-length fusions were constructed as follows. A PCR fragment containing *whiA* N-terminus domain, C-terminus domain and full-length were amplified with oligonucleotide pairs LB11-LB12, LB13-LB14 and LB11-LB14, respectively. Genomic DNA of strain 168 was used as template. *BamHI* and *EcoRI* restriction sites, a flexible linker and terminator were inserted into the primers. Each PCR product and the *amyE-* integration vector pHJS105 (55) were digested with *BamHI* and *EcoRI* restriction enzymes and ligated. The resulting plasmids were named pLB19, pLB20 and pLB18, respectively, verified by sequencing and transformed into *B. subtilis* 168 and competent cells, resulting in strains LB230, LB231 and LB232, respectively. Each strain was transformed with genomic DNA from *whiA* knockout KS400 (7), resulting in strains LB295, LB296 and LB294, respectively. The cellular localization of the GFP fusion proteins was determined using fluorescence microscopy.

### Yeast-two hybrid assays

Proteins of interest were expressed in *Saccharomyces cerevisiae* strain PJ69-4a as fusions to the GAL4 binding domain BD or activating domain AD, from the vectors pGBDU-C1 and pGAD-C1, respectively. The *whiA* N-terminal and C-terminal domains were cloned into a pGBDU bait vector by gap repair and directly transformed into *S. cerevisiae* strain PJ69-4a. The DNA sequences of all cloned fragments were verified by sequencing. These baits were used to screen a *B. subtilis* prey library essentially as previously described (9). In brief, three *B. subtilis* genomic libraries were constructed in *E. coli*, restrictions of the 4.2 Megabase *B. subtilis* chromosome produced approximately 1.6 x 10^5^ DNA ends that could be ligated into the pGAD prey vectors. Each library contained at least 2.5 x 10^6^ clones, thus providing a 15-fold redundancy. The PJ69-4a yeast strain was transformed by each library DNA and at least 1.5 x 10^7^ prey-containing colonies were harvested and pooled. The library-containing cells were mated with bait-containing cells. The mixture was plated on rich YEPD medium and incubated for 5 h at 30 °C. Cells were collected, washed and spread on synthetic complete medium plates lacking the amino acids leucine and histidine and the nucleotide uracil (SC-LUH). To calculate the mating efficiencies and the number of diploids, cells were also spread on SC-L and SC-LU plates. A screening is covered if the number of diploids is greater than 1 x 10^6^ and the mating efficiency greater than 20 %. After 10-12 days of incubation at 30 °C, the colonies obtained were transferred to the SC-LUA (synthetic complete medium lacking leucine, uracil and adenine) and SC-LUH medium and incubated for 3-5 days. The interaction candidates were identified by PCR amplification and sequencing of the DNA inserts in the prey plasmids. To screen for false-positive interactions, protein-encoding prey plasmids were rescued from His^+^ Ade^+^ colonies, reintroduced in PJ69–4alfa strain by transformation and subjected to a mating with cells containing: *i*) an empty bait vector, *ii*) the initial bait used in the screen and *iii*) a variety of unrelated baits. The diploid cells were tested for expression of the interaction phenotypes (His^+^ and Ade^+^). Specific interactions were reproducible with the initial bait and not associated with self-activation or stickiness of the prey protein. The interactions not fulfilling these criteria corresponded to false positives and were discarded.

### Chromosome capture by Hi-C

Cultures were grown in LB media with shaking and samples for Hi-C were collected at exponential growth phase (OD_600_ 0.6). Hi-C was carried out exactly as described before with digestion using *HindIII* (56). Hi-C matrices were constructed using the Galaxy HiCExplorer webserver (57). Briefly, paired-end reads were mapped separately to the *B. subtilis* genome (NCBI Reference Sequence NC_000964.3) using very sensitive local setting mode in Bowtie2 (Galaxy v.2.3.4.2). The mapped files were used to build the contact matrix using the tool hicBuildMatrix (Galaxy v.2.1.2.0) using a bin size of 10 kb, and *HindIII* restriction site (AAGCTT) and AGCT as the dangling sequence. The contact matrix was then used for further analysis and visualization using the interactive browser-based visualization tool ‘Bekvaem’ essentially as described before (56).

### Microscopy

Exponentially growing cells were stained with the fluorescent membrane dye FM-95 and the DNA was stained with DAPI. Cells were grown overnight on LB agar plates. A single colony was streaked out on LB agar plates supplemented with 0.1 % xylose for the induction of expression, grown for ~6 h and subsequently mounted on microscope slides covered with a thin film of 1 % agarose. Microscopy was performed on an inverted fluorescence Nikon Eclipse Ti microscope. The digital images were acquired and analysed with ImageJ v.1.48d5 (National Institutes of Health).

### Metabolome analysis

The main culture (20 ml) was inoculated with an exponentially growing overnight culture to an initial OD_500_ of 0.05. The optical density was monitored and 2 ml cell suspension was sampled. Three experiments were carried out to provide the necessary biological replicates. During cultivation, the pH value was determined at each sampling time point by using HI 2211 pH/mV/uC bench meter (Hanna instruments Deutschland GmbH, Kehl, Germany). 2 ml of cell culture medium were taken at 60, 120, 180, 240, 300, 360, 420 and 480 min by sterile filtration, using a 0.45 mm pore size filter (Sarstedt AG, Nuernberg, Germany), to get sterile extracellular metabolite samples of the bacterial culture, and directly frozen until measurement. ^1^H-NMR analysis was carried out as described previously (58). In brief, 400 μl of the sample was mixed with 200 μl of a sodium hydrogen phosphate buffer (0.2 M, pH 7.0) to avoid chemical shifts due to pH, which was made up with 50 % D_2_O. The buffer also contained 1 mM trimethylsilyl propanoic acid-d_4_ (TSP) which was used for quantification and also as a reference signal at 0.0 ppm. To obtain NMR spectra, a 1D-NOESY pulse sequence was used with a presaturation on the residual HDO signal. A total of 64 FID scans were performed with 600.27 MHz and at a temperature of 310 K using a Bruker AVANCE-II 600 NMR spectrometer operated by TOPSPIN 3.1 software (both from Bruker Biospin). For qualitative and quantitative data analysis, we used AMIX (Bruker Biospin, version 3.9.14). We used the AMIX Underground Removal Tool on obtained NMR-spectra to correct the baseline, thereby using the following parameters: left border region 20 ppm and right border region −20 ppm and a filter width of 10 Hz. Absolute quantification was performed as previously described (58). In brief, a signal of the metabolite, either a complete signal or a proportion, was chosen manually and integrated. The area was further normalized on the area of the internal standard TSP and on the corresponding number of protons and the sample volume. For statistical comparison of extracellular metabolite data and growth, barcharts, and XY-plots, we used Prism (version 6.01; GraphPad Software). The time-resolved extracellular metabolite concentrations were log_2_ (*x* + 1) transformed for the separation via PCA. The PCA was done using PAST v3.16 with auto-scaled data (59).

### Transcriptome analysis

Cells (2 ml cultures) were spun down (30 s Eppendorf centrifuge, 14,000 rpm, 4 °C), resuspended in 0.4 ml ice-cold growth medium and added to a screw cap Eppendorf tube containing 1.5 g glass beads (0.1 mm), 500 μl phenol:chloroform:isoamyl alcohol (25:24:1), 50 μl 10 % SDS and 50 μl RNAse free water (60). All solutions were prepared with diethylpyrocarbonate (DEPC)-treated water. After vortexing, tubes were frozen in liquid nitrogen and stored at −80°C. Cells were broken using a bead-beater for 4 min at room temperature. After centrifugation, the water phase was transferred to a clean tube containing 400 μl chloroform, after vortexing and centrifugation, the water phase was used for RNA isolation with High Pure RNA Isolation Kit (Roche Diagnostics GmbH, Mannheim, Germany), yielding >3 μg total RNA per sample. TapeStation System (Agilent) was used for checking the integrity of the RNA, and RIN values of 8.3 – 9.2 were obtained. For next-generation sequencing, a ribosomal RNA depletion was performed on the total RNA using the Ribo-Zero rRNA Removal Kit (Gram-Positive Bacteria) (Illumina). Bar-coded RNA libraries were generated according to the manufacturers’ protocols using the Ion Total RNA-Seq Kit v2 and the Ion Xpress RNA-Seq barcoding kit (Thermo Fisher Scientific). The size distribution and yield of the barcoded libraries were assessed using the 2200 TapeStation System with Agilent D1000 ScreenTapes (Agilent Technologies). Sequencing templates were prepared on the Ion Chef System using the Ion PI Hi-Q Chef Kit (Thermo Fisher Scientific). Sequencing was performed on an Ion Proton System using an Ion PI v3 chip (Thermo Fisher Scientific) according to the instructions of the manufacturer. After quality control and trimming the sequence reads were mapped onto the genome (genome-build-accession NCBI Assembly: GCA_000009045.1) using the Torrent Mapping Alignment Program. The Ion Proton system generates sequence reads of variable lengths, and this program combines a short read algorithm (61), and long read algorithms (62) in a multistage mapping approach. The gene expression levels were quantified using HTseq (63). The data was normalized and analysed for differential expression using R statistical software and the DESeq2 package (64). The RNA-seq data have been submitted to and are accessible in the Gene Expression Omnibus (GEO) using accession number GSE121479.

### Lipid analysis

The fatty acid composition was determined from cells growing in Amber medium when the cultures reached an OD_600_ of approximately 0.5. Cells were harvest by centrifugation at 10.000x rcf for 5 min at 4 °C, washed once with 0.9 % ice-cold NaCl, and submitted to flash freeze in liquid N2. Fatty acids were analyzed as fatty acid methyl esters (FAME) using gas chromatography. All analyses were carried out in triplicates at theLaboratory Genetic Metabolic Disease, Amsterdam UMC.

### Laurdan GP spectroscopy

For the measurement of membrane fluidity in batch cultures as reported before (49), cells were grown in either LB or Spizizen minimal salt medium (SMM) to an OD of approximately 0.5, followed by 5 min incubation with 10 mM Laurdan. Subsequently, cells were washed three times with pre-warmed buffer containing 50 mM Na_2_HPO_4_/NaH_2_PO_4_ pH 7.4, 0.1 % glucose and 150 mM NaCl with and without the membrane fluidizer benzyl alcohol (30 mM). The Laurdan fluorescence intensities were measured at 435±5 nm and 490±5 nm upon excitation at 350±10 nm, using a Tecan Infinite 200M fluorometer. The Laurdan generalized polarization (GP) was calculated using the formula GP = (I_435_ – I_490_) / (I_435_ + I_490_).

## ACKNOWLEDGEMENTS

We would like to thank Henrik Strahl (Newcastle University) for scientific assistance with the lipid analysis and for insightful discussions, Johan Westerhuis (University of Amsterdam) for his insights about the analysis of the exometabolome, Selina van Leeuwen (MAD, UvA) for providing excellent sequencing services, and to all the members of the Bacterial Cell Biology group (University of Amsterdam), and specially to all the members of the Advanced Multidisciplinary Training in Molecular Bacteriology (AMBER) EU Marie Curie Initial Training Network (ITN). The research was funded by EU Marie Curie ITN grant AMBER (317338), Marie Curie CIG grant DIVANTI (618452), European Commission MCSA-IF 749510, and STW Vici grant 12128.

